# Peripheral *Htt* silencing does not ameliorate central signs of disease in the B6. *Htt^Q111/+^* mouse model of Huntington’s Disease

**DOI:** 10.1101/096990

**Authors:** Sydney R. Coffey, Robert M. Bragg, Minnig Shawn, Seth A. Ament, Glickenhaus Anne, Shelnut Daniel, José M. Carrillo, Dominic D. Shuttleworth, Rodier Julie-Anne, Noguchi Kimihiro, C. Frank Bennett, Nathan D. Price, Holly B. Kordasiewicz, Jeffrey B. Carroll

## Abstract

Huntington’s disease (HD) is an autosomal dominant neurodegenerative disease whose neuropathological signature is a selective loss of medium spiny neurons in the striatum. Despite this selective neuropathology, the mutant protein (huntingtin) is found in virtually every cell so far studied, and, consequently, phenotypes are observed in a wide range of organ systems both inside and outside the central nervous system. We, and others, have suggested that peripheral dysfunction could contribute to the rate of progression of striatal phenotypes of HD. To test this hypothesis, we lowered levels of huntingtin by treating mice with antisense oligonucleotides (ASOs) targeting the murine *Huntingtin* gene. To study the relationship between peripheral huntingtin levels and striatal HD phenotypes, we utilized a knock-in model of the human HD mutation (the B6.*Htt^Q111/+^* mouse). We treated mice with ASOs from 2-10 months of age, a time period over which significant HD-relevant signs progressively develop in the brains of *Htt^Q111+^* mice. Peripheral treatment with ASOs led to persistent reduction of huntingtin protein in peripheral organs, including liver, brown and white adipose tissues. This reduction was not associated with alterations in the severity of HD-relevant signs in the striatum of *Htt^Q111/+^* mice at the end of the study, including transcriptional dysregulation, the accumulation of neuronal intranuclear inclusions, and behavioral changes such as subtle hypoactivity and reduced exploratory drive. These results suggest that the amount of peripheral reduction achieved in the current study does not significantly impact the progression of HD-relevant signs in the central nervous system.

## Introduction

Huntington’s disease (HD) is an autosomal dominant neurodegenerative disorder caused by a glutamine-encoding CAG expansion near the 5’ end of the Htt gene [1]. The symptoms of HD are progressive cognitive, affective and characteristic motor symptoms, generally beginning in mid-life and progressing inexorably to death approximately 15 years after initial symptoms present [2]. These symptoms have been linked to progressive dysfunction and cell death in corticostriatal circuits in mutation carriers [3], though atrophy in widespread brain regions are also implicated to varying degrees [4]. By late stages of the disease, nearly complete loss of striatal projection cells has occurred [5].

Thanks to HD’s complete penetrance, and the wider interest in the pre-symptomatic progression of neurodegenerative diseases, HD mutation carriers are amongst the most robustly studied populations in medicine. Asymptomatic HD mutation carriers progressive phenotypes have been characterized in many dozens of studies, in some cases for longer than 10 years of continuous observation [6,7]. These observational studies reveal widespread peripheral signs associated with carrying the HD mutation, in addition to progressive neurological symptoms [8]. The appearance of peripheral phenotypes may not be surprising, given that the Huntingtin protein (HTT) and transcript (*Htt*) are widely, and consistently, expressed in every cell type so far studied [9]. Observed peripheral changes in HD mutation carriers include: subtly enhanced immune activation [10], progressive reductions in hepatic mitochondrial function [11,12], and progressive loss of lower limb strength [13]. We have proposed [9] that these symptoms are worth understanding for several reasons: first - peripheral dysfunction may contribute to CNS pathology directly; second – they may autonomously lead to patient morbidity and/or mortality; and, finally - they may uncover novel aspects of HTT function by revealing physiological pathways impacted by its mutation in other organs.

We set out to test the first of these hypotheses by peripherally silencing HTT in the B6.*Htt^Q111/+^* mouse model of the HD mutation, which precisely recapitulate the genetics of human HD, expressing a single mutant allele from the endogenous murine Htt locus [14]. Compared to HD patients and transgenic mouse models, these mice have relatively subtle signs of disease but do present with a wide range of molecular changes, especially in the striatum, the most vulnerable region of the brain in HD [15],[16]. To study the relationship between peripheral huntingtin levels and disease, we silenced huntingtin (and mutant huntingtin) using peripherally-restricted antisense oligonucleotides (ASOs). ASOs are short strands of chemically-modified deoxyribonucleotides which hybridize with target mRNA and modulate its processing in various ways, including RNAseH-mediated degradation [17]. Because they are large charged molecules, peripherally-administered ASOs are effectively excluded from the brain by the blood brain barrier [18]. We took advantage of this selective localization to investigate whether peripheral silencing of HTT would impact signs of the disease in the central nervous system (CNS). We were particularly interested in the impact of hepatic silencing of *Htt* on HD symptoms, because the liver is an important nexus for brain-body cross-talk. The liver synthesizes glucose (via gluconeogenesis) and ketone bodies (via ketogenesis), which serve as critical substrates for the brain between meals and while fasting [19]. The liver also regulates whole-body levels of nitrogenous waste products, including urea, a critical function for the preservation of brain health. Increased brain urea levels have been reported in human HD patients and model sheep [19,20]. Likewise, increased circulating ammonia has been reported in mouse models of disease [21]. Generally, liver failure is associated with neurological sequelae, hepatic encephalopathy, which, like HD, involves dysfunction in corticostriatal circuits, increased inflammation and excitotoxicity [22].

Peripheral interventions have been successful at relieving central signs of disease in other models of genetic neurodegenerative disease. For example, peripheral silencing of polyglutamine expanded androgen receptor in mouse models of spinal muscular bulbar atrophy leads to robust rescue of a number of phenotypes, including motor improvement and survival extension [23]. Similarly, peripherally administration of ASOs guiding enhanced exon skipping in the transcript for survival motor neuron 2 gene leads to dramatic improvement of phenotypes in mouse models of spinal muscular atrophy [23]. In HD, protecting mice from urea cycle dysfunction via a low protein diet is sufficient to improve CNS-resident signs of HD [24]. In Parkinson’s Disease (PD), fibrillar alpha-synuclein has been proposed to travel from neurons in the enteric nervous system, via the vagus, to the brainstem [25]. This mechanism is supported by the markedly reduced risk of sporadic PD in patients undergoing vagotomy [26], and the ability of fibrillar alpha-synuclein to travel from the gut to the brainstem of experimental animals [27]. Similar prion like spreading has been proposed in HD - transplantation of human HD fibroblasts into the ventricles of wild-type animals results in the formation of mutant Huntingtin aggregates in wild-type cells [28]. In short, peripheral silencing of neurodegenerative disease proteins has the potential to impact disease progression.

Given the important links between peripheral organ function and brain health, and evidence that the HD mutation is associated with alterations in whole-body physiology, we tested the relationship between peripheral huntingtin silencing and striatal signs of HD in the B6.*Htt^Q111/+^* model of HD. Using systemically-delivered ASOs, we silenced hepatic HTT during a window in which a range of progressive striatal signs of disease develop. This treatment robustly reduced HTT levels in liver and adipose tissues, but did not alter striatal signs of HD, suggesting these are independent of hepatic dysfunction in this model.

## Results

### HTT knockdown

Beginning at 2 months of age, we intraperitoneally (IP) injected *Htt^Q111/+^* and wild-type littermates with *Htt*-targeted ASO to suppress both endogenous and expanded HTT (Fig 1A). Weekly deliveries of 50 mg per kg body mass (mpk) of either an ASO targeting *Htt* (hereafter ‘*Htt* ASO’), an off-target control ASO (hereafter ‘control ASO’), or saline alone continued until 10 months of age, at which point mice were sacrificed. Peripheral HTT knockdown by *Htt* ASO treatment was confirmed in three tissues of interest - liver, perigonadal white adipose tissue, and interscapular brown adipose tissue - using mesoscale discovery (MSD) assays, which quantify total and polyglutamine expanded HTT (Fig 1B) [29]. Compared to control ASO-treated mice, *Htt* ASO-treated mice had significantly reduced HTT levels in all peripheral tissues examined (effect of treatment in the liver: *F_(1, 15)_* = 79.6, *p* = 2.2 × 10^−7^, white adipose tissue: *F_(1, 15)_ = 39.6, p* = 1.4 × 10^−5^, and brown adipose tissue: *F_(1, 15)_ =89.2, p* = 1.1 × 10^−7^). To confirm the non-allele-specificity of the chosen *Htt* ASO, mutant HTT (mHTT) levels were independently determined using an antibody pair specific for mutant huntingtin. Consistent with the lack of sequence difference between wild-type and mutant *Htt* at the ASO target sequence, similar patterns of suppression of mHTT are observed in all peripheral tissues tested (effect of treatment in the liver: *F_(1, 15)_ = 163.2, p* = 1.8 × 10^−9^, white adipose tissue: *F_(1, 15)_ = 53.2, p* = 2.7 × 10^−6^, and brown adipose tissue: *F_(1, 15)_ = 233.3, p* = 1.5 × 10^−10^). In contrast to its effects in peripheral tissues, intraperitoneal injection with *Htt* ASO-treatment had no effect on total HTT (*F*_(2, 18)_=0.73, *p* = 0.49) or mutant HTT (*F_(2, 9)_ = 0.16, p* = 0.86) levels in the striatum (Fig 1C-D). Amongst examined tissues, ASO-mediated suppression of total HTT was most efficient in the white adipose tissue (71% knockdown), and least efficient in the liver (65% knockdown). These data confirm that we were able to robustly silence HTT in peripheral organs (liver, white adipose and brown adipose) without reducing levels in the striatum. To verify HTT levels did not fluctuate throughout the trial, we examined two interim silencing cohorts, which were sacrificed prior to study completion (Fig 1A). We selected two timepoints approximately one-and two-thirds through the 8-month trial and quantified HTT levels using the MSD assay. Interim results were consistent with the endpoint analysis described above (Figure in S3 Figure). As a qualitative assessment of *Htt* ASO uptake in other peripheral tissues, we stained fixed organs with an antibody reactive to the ASO backbone. We find that *Htt* ASO uptake appears most efficient in the liver, spleen, and kidney, with modest uptake in the heart and skeletal muscle (Figure in S4 Figure).

**Fig 1.**
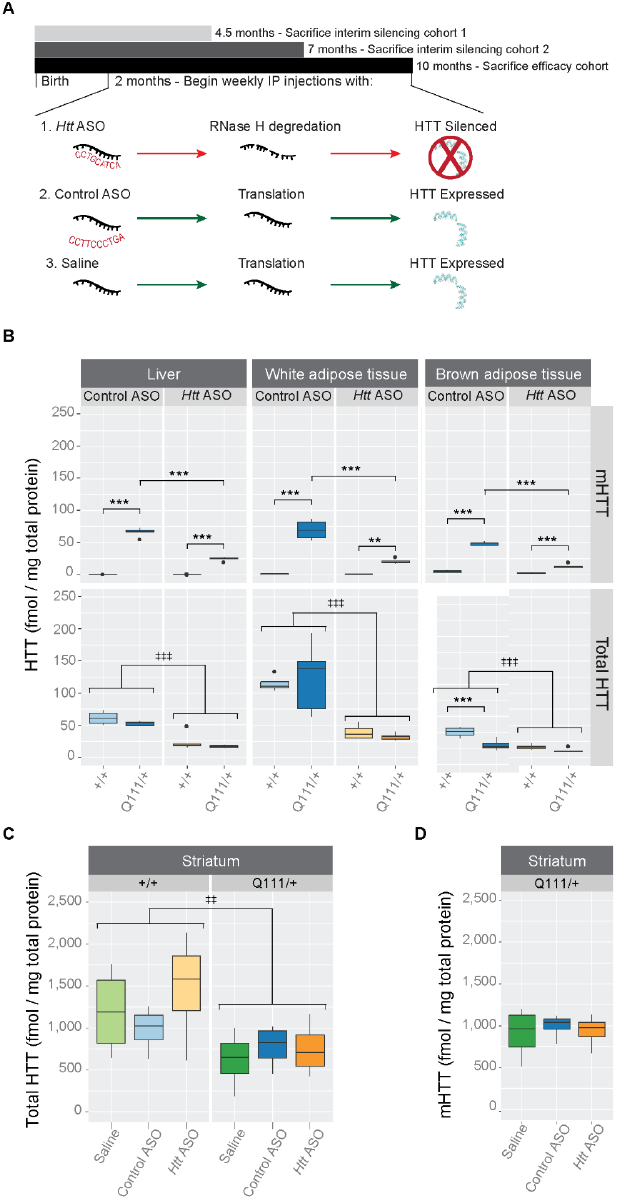
IP-delivered *Htt* ASO suppresses both wild-type and mutant huntingtin in the periphery, but not striatum. (A) Starting at 2 months of age and continuing until 10-months, *Htt^Q111/+^* or *Htt^+/+^* mice received weekly IP injections of *Htt* ASO, control ASO, or saline alone. In parallel to this efficacy cohort, two interim silencing cohorts were established to verify continuous suppression of HTT by *Htt* ASO. (B) Levels of total and mutant HTT were quantified by MSD assay in three peripheral tissues: liver, white and brown adipose tissue. *Htt* ASO treatment significantly reduced levels HTT compared to control ASO treatment in all three tissues. (C-D) Treatment with *Htt* ASO did not alter striatal levels of total (C) or mutant (D) huntingtin. All data are presented as boxplots. * p ≤ 0.05, ** p ≤ 0.01, *** p ≤ 0.001: by Tukey’s HSD pairwise comparisons ‡ p ≤ 0.05, ‡‡ p ≤ 0.01, ‡‡‡ p ≤ 0.001: by factorial ANOVA

## Central pathological signs

Striatal-specific accumulation of neuronal intranuclear inclusions (NII’s) is a defining feature of the aging *Htt^Q111/+^* mouse [15]. We therefore considered whether peripheral suppression of HTT affected central aggregate load by counting aggregates immunoreactive for p62/Sqstm-1, a receptor for cargos destined to be degraded by selective macroautophagy [30]. In order to restrict our analysis to neurons, we created a mask for our images by selecting only cells immunoreactive for Rbfox3 (also known as NeuN), a pan-neuronal marker protein [31]. Consistent with previous investigations [32], we observe accumulation of p62-immunoreactive NII’s in striatal neurons from *Htt^Q111/+^* mice (22.5 ± 0.14% of *Htt^Q111/+^* striatal neurons have NIIs (0.5 - 5 μm^2^ size cutoff) compared to 0 ± 0.3% of *Htt+/+* neurons, Fig 2). Peripheral treatment of *Htt^Q111/+^* mice with *Htt* ASOs does not impact this phenotype (effect of treatment: F_(2,24)_= 1.6, p = 0.2, Fig 2).

**Fig 2.**
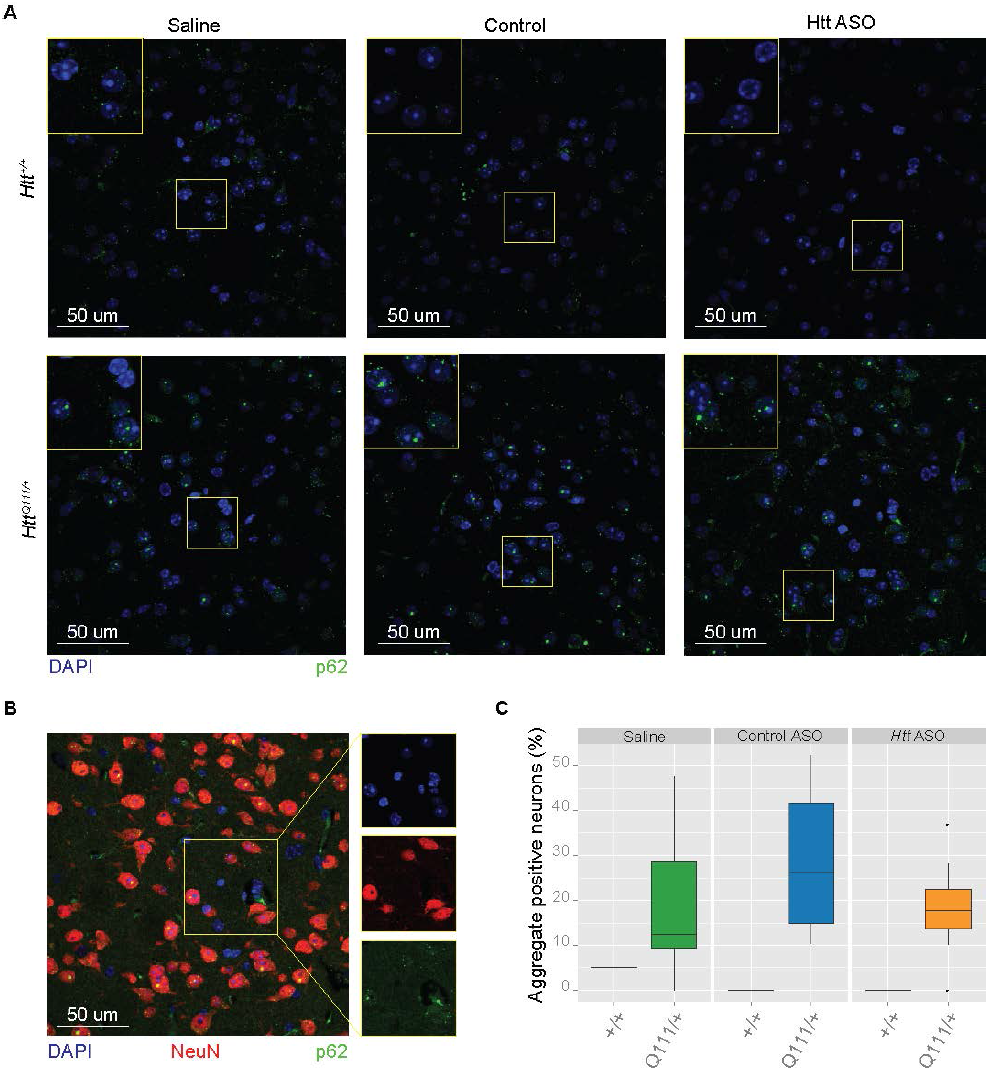
Peripheral silencing of *Htt* does not alter p62-immunoreactive neuronal intranuclear inclusions in the dorsolateral striatum. (A) Representative images show nuclei (DAPI, blue) and autophagy adapter p62 (green) for each genotype and treatment condition. For clarity, staining for neuronal marker NeuN was omitted in (A). However, a neuronal (NeuN) mask was used to selectively quantify p62 aggregates in neurons, therefore a representative image of an *Htt^Q111/+^* mouse is shown in (B). Quantification of staining among *Htt^Q111/+^* mice (C) reveals no differences in the percent of neurons with intranuclear inclusions between treatment groups.

Progressive, striatal-specific, transcriptional dysregulation is another feature of HD mouse models, including the *Htt^Q111/+^* mouse [15,16]. We therefore examined striatal transcriptional dysregulation using both targeted (QRT-PCR) and untargeted (RNA Sequencing, RNASeq) methods. As expected, 10-month-old *Htt^Q111/+^* mice have reduced levels of key striatal synaptic and signaling genes, including *Drd1a* and *Pde10a* (Fig 3). In each case, transcript levels were reduced in the striatum of *Htt^Q111/+^* mice relative to *Htt^+/+^* mice using both RNASeq (*Drd1a*: F_(1,29)_ = 103.78, FDR =1.66 × 10^−12^, *Pde10a*: F_(1,29)_ = 168.06, FDR = 5.94 × 10^×15^) and QRT-PCR detection methods (*Drd1a*: F_(1,53)_ = 67.97, p = 4.57 × 10^−11^, *Pde10a*: F_(1,53)_ = 44.47, p = 1.54 × 10^−8^). To ensure neither our QRT-PCR or RNAseq methods preferentially detected only downregulated genes, we quantified HD-relevant transcripts previously reported as upregulated in the striatum of *Htt^Q111/+^* mice [15]. For instance, we successfully replicated increased levels of *Islr2* and *N4bp2* in 10-month *Htt^Q111/+^* mice (Fig 3) using both RNAseq (*Islr2*: F_(1,29)_ = 42.84, FDR = 3.45 × 10^−8^ and *N4bp2*: F_(1,29)_ = 10.96, FDR = 1.67 × 10^−3^) and QRT-PCR (*Islr2*: F_(1,52)_ = 10.65, p = 1.95 × 10^−3^ and *N4bp2*: F_(1,53)_ = 7.68, p = 7.68 × 10^−3^). Peripheral treatment with *Htt* ASO did not impact expression level of any of these transcripts, suggesting that striatal transcriptional dysregulation was not rescued by this treatment.

**Fig 3.**
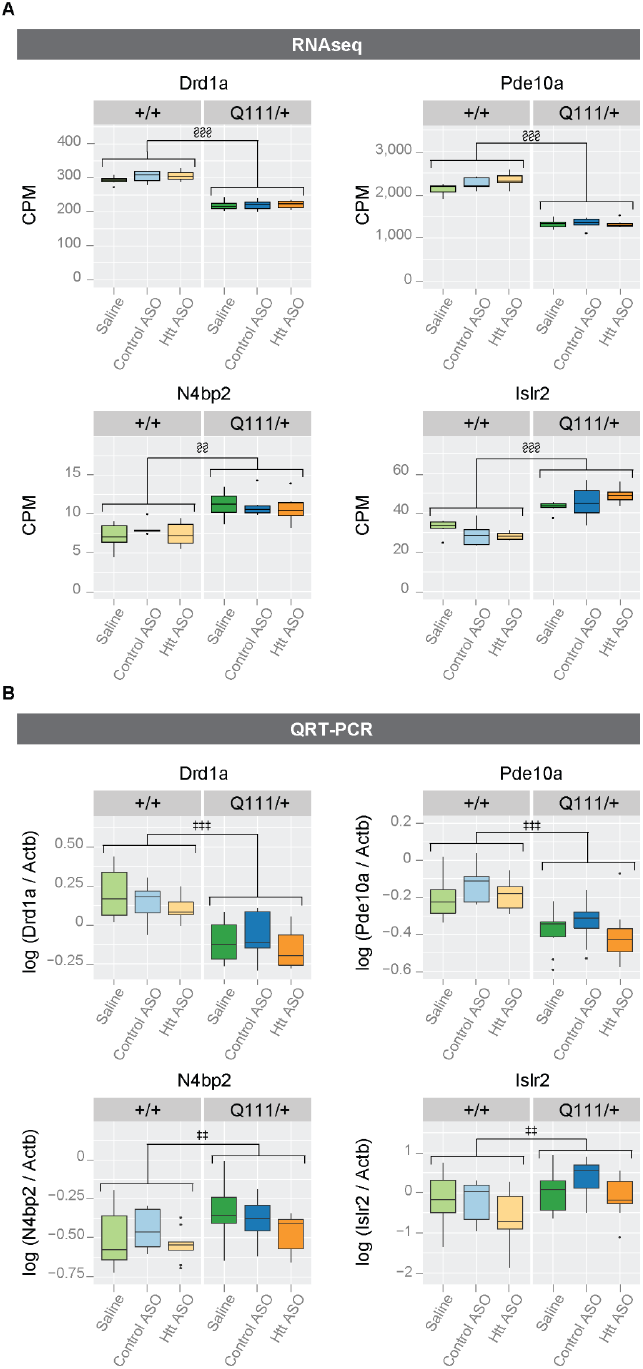
Peripheral *Htt* silencing does not rescue striatal transcriptional dysregulation in *Htt^Q111/+^* mice. Using both RNAseq (A) and QRT-PCR (B), we assessed transcriptional markers of HD-like pathology in the striatum. Four transcripts, *Drd1a*, *Pde10a*, *N4bp2*, and *Islr2* were selected to illustrate the agreement between QRT-PCR and RNAseq results. We successfully replicated HD-relevant transcriptional changes previously characterized in *Htt^Q111/+^* and wild-type mice, but *Htt* ASO treatment failed to rescue these phenotypes. Abbreviations - CPM: counts per million ‡ p ≤ 0.05, ‡‡ p ≤ 0.01, ‡‡‡ p ≤ 0.001: by factorial ANOVA § FDR ≤ 0.05, §§ FDR ≤ 0.01, §§§ FDR ≤0.001

## Behavior and Body Weight

One month prior to sacrifice, we investigated whether peripheral ASO treatment improved any HD-relevant behavioral changes in the *Htt^Q111/+^* mouse. We first examined exploratory behavior in an open field task, testing mice during the first 8 hours of their dark cycle in an illuminated room maintained at approximately 475 lux. Consistent with previous investigations [33], exploratory activity in a 10 minute open field task was modestly reduced in *Htt^Q111/+^* mice, compared to *Htt_+/+_* mice (Fig 4A; 8.95% reduction, effect of genotype, *F*_(1,93)_ = 6.6, *p* = 0.01). There was no main effect of treatment (*F*_(2,93)_ = 0.3, *p* = 0.7) or genotype/treatment interaction (*F*_(2,93)_ = 0.3, *p* = 0.7), suggesting that peripheral *Htt*-ASO treatment was not able to improve the mildly hypoactive phenotype observed in HD mice. To determine if the reduction in exploratory activity observed in HD mice may be caused by subtle motor deficit, we calculated the average velocity of each mouse and observe a mild reduction in *Htt^Q111/+^* mice (4.9% reduction; *F*_(1,93)_ = 5.7, *p* = 0.02), but no effect of treatment (*F*_(2,93)_ = 0.5, *p* = 0.6) or genotype/treatment interaction (*F*_(2,93)_ = 0.2, *p* = 0.86). We also quantified thigmotaxis, the tendency to explore the outer walls of the open field arena compared to the center, as a proxy for anxiety levels [34]. We observed a modest increase in thigmotaxis in *Htt^Q111/+^* mice (Fig 4B; 8.3% increase; genotype, *F*_(1,93)_ = 6.3, *p* = 0.01), but no effect of treatment (*F*_(2,93)_ = 1.9, *p* =0.2) or genotype/treatment interaction (*F*_(2,93)_ = 0.6, *p* = 0.6). These results suggest that 9-month-old *Htt^Q111/+^* mice are mildly hypoactive and potentially anxious, but that reducing peripheral HTT levels does not improve these phenotypes.

**Fig 4.**
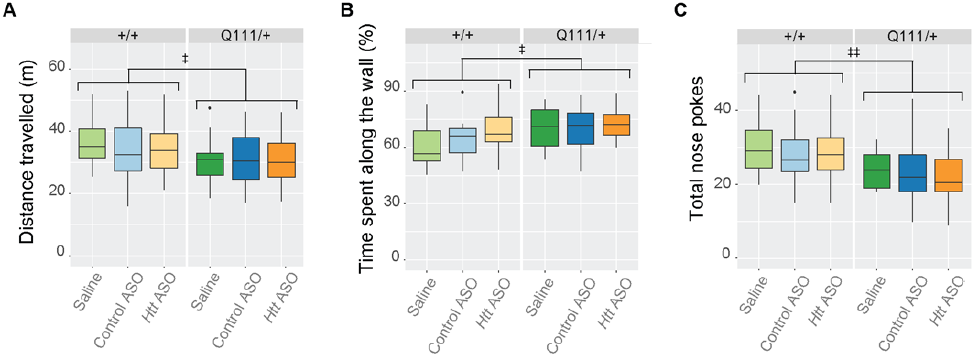
Hypoactivity and increased anxiety-like phenotypes are not improved by peripheral *Htt* silencing. During open field exploration, *Htt^Q111/+^* mice travel less distance than *Htt^+/+^* mice (A) and spend more time along the outer walls of the arena (B). This thigmotactic tendency combined with observations that *Htt^Q111/+^* mice initiate fewer total investigations of both objects during the novel object location task (C) suggest that *Htt^Q111/+^* mice exhibit anxiety-like phenotypes and are less motivated to explore. However, *Htt* ASO treatment failed to modulate these phenotypes.‡ p ≤ 0.05, ‡‡ p ≤ 0.01, ‡‡‡ p ≤ 0.001: by factorial ANOVA

We next examined long term spatial memory in *Htt^Q111/+^* mice using a novel object location task (OLT), which exploits the spontaneous tendency of rodents to preferentially explore an object that has been displaced to an unfamiliar location following a delay period, in this case 24 hours. In contrast to previous reports of impairments in small cohorts of 6-month old *Htt^Q111/+^* mice on this task [35], we find both *Htt^+/+^* and *Htt^Q111/+^* mice have a preference for the novel object (group average of 58.2 ±12.3% of total nose pokes were towards the novel object; test for preference for novel object location: *Htt^+/+^:* □^*2*^ = 50.4*, p* <.001; *Htt*^*Q111/+*^: □^*2*^ = 23.1, *p* <.001). A 2×3 between subjects ANOVA revealed no main effect of genotype, *F*_(1,92)_ = 0.2, *p* = 0.7, no main effect of treatment, *F*_(2,92)_ = 0.8, *p* = 0.5, and no interaction between genotype and treatment *F*_(2,92)_ = 2.2, *p* = 0.1, indicating that *Htt^Q111/+^* mice display intact spatial long-term memory at 9 months of age (Figure in S6 Figure). Interestingly, a 2×3 between subjects ANOVA revealed a main effect of genotype (Fig 4C; *F*_(1,92)_ = 14.15, *p* < 0.01), such that *Htt^Q111/+^* mice conducted significantly fewer investigations of either object during the novel object location task (19.1% reductions in total investigations). These reductions suggest that previous reports of altered long-term memory could be confounded by neophobia or motivational changes in *Htt^Q111/+^* mice at this age, rather than spatial memory deficits. There was no main effect of *Htt* ASO treatment (*F*_(2,92)_ = 0.5, *p* = 0.6) and no interaction between genotype and treatment (*F*_(2,92)_ = 0.3, *p* = 0.8), suggesting that ASO treatment was unable to rescue this altered behavior in *Htt^Q111/+^* mice.

Throughout the study we observed body mass in order to track any toxic liabilities of ASO treatment. We used a minP-based parametric bootstrap multiple comparison procedure [36],[37] to assess differences in body mass throughout the 35 weeks of treatment. The minP procedure provides adjusted *p*-values that correctly account for the familywise error rate as the error control criterion based on the raw *p*-values that are calculated from Welch’s *t*-test. As the *p*-value adjustment requires the knowledge of the underlying distribution of the minimum of *p*-values under the null hypothesis of no weight difference, we approximated the distribution using the parametric bootstrap with 10,000 bootstrap resamples from the mean-zero multivariate normal distribution with the estimated covariance matrix [37]. Because the raw *p*-values are used as the test statistics in the minP procedure, we report the minimum of the raw *p*-values (*P*_min_) as the test statistic for the hypothesis of interest, and the adjusted *p*-value as the corresponding *p*-value.

Consistent with previous observations [15,33], we did not observe changes in body weight in *Htt^Q111/+^* mice relative to wild-types (*P*_min_ = 0.27, *p* = 0.92). Therefore, we solely focused on the treatment effects by pooling weight observations from both the *Htt^Q111/+^* and *Htt^+/+^* mice. As we were interested in the mean weight difference (if any) between saline and control ASO treated mice, as well as the mean weight difference between the grouped saline and control ASO treated mice compared to *Htt* ASO treated mice, these two hypotheses were tested simultaneously using the minP-based parametric bootstrap multiple comparison procedure. After 30 weeks of *Htt* ASO treatment, mice exhibit modestly decreased body mass (Fig 5B), weighing 5% less than grouped control and saline treated mice (*P*_min_ <.001, *p* = 0.0001). This is clearly exhibited in Fig 5A, where the adjusted *p*-values are less than 0.05 after 30 weeks. On the other hand, there was no significant difference at any week between the saline and control-ASO treated mice (*P*_min_ = 0.04, *p* = 0.58).

**Fig 5.**
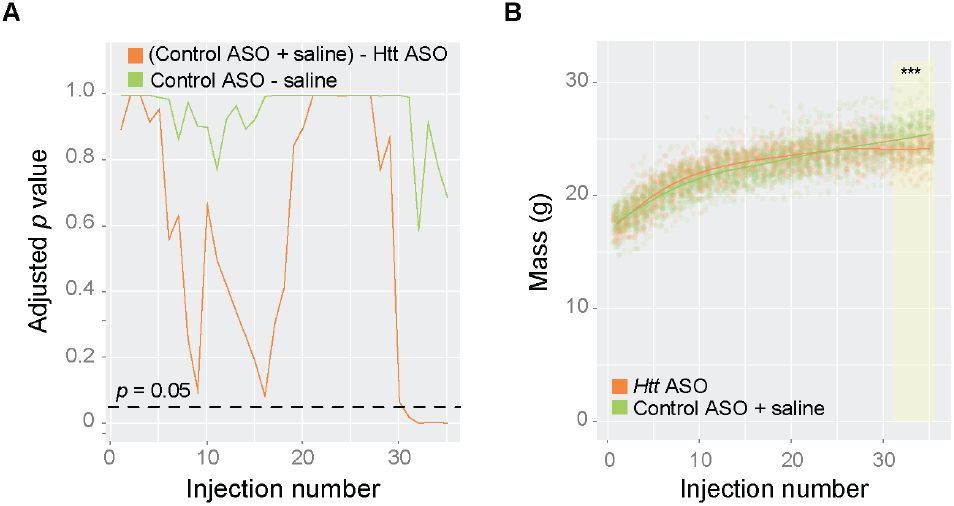
Prolonged treatment with *Htt* ASO leads to modest reductions in body weight. Body mass was recorded weekly during IP injections and examined longitudinally using minP-based parametric bootstrap multiple comparison procedure [36,37]. (A) No significant differences were observed between control ASO and saline treated mice (*p* = 0.58); therefore, control ASO and saline treated mice were grouped and compared to *Htt* ASO treated mice. (B) After 30 injections, the body mass of *Htt* ASO treated mice was decreased by 5% (denoted by highlighted region; overall curve comparison p = 0.0001) compared to grouped control ASO and saline treated mice.

## Discussion

Peripheral phenotypes are widely observed in HD mutation carriers, thanks to the ubiquitous expression of the mutant *Htt* allele. We have suggested that peripheral dysfunction could potentially directly contribute to the progression of signs of HD in the CNS [9], a hypothesis we tested here by silencing *Htt* in peripheral organs. We find that effective silencing of hepatic and adipose *Htt* during a window in which the progression of central signs occurs rapidly [15] does not impact the progression of signs of disease in the CNS. These symptoms include transcriptional dysregulation, accumulation of neuronal intranuclear inclusions, and HD-relevant behavioral changes.

Our experiments were designed to test whether direct reduction of mutant huntingtin in peripheral organs, rather than than correction of downstream physiological perturbations, is generally beneficial for the CNS. Several previous studies in HD mouse models have examined the effect of peripherally-restricted interventions on the progression of some CNS-resident phenotypes. For example, imposition of a low-protein diet reduces circulating ammonia and citrulline levels and is associated with improved behavioral and CNS pathological signs of HD in both knock-in and transgenic mouse models [21]. Cross-genotype bone marrow transplants dampen peripheral immune activation in transgenic BACHD mice, and are associated with preservation of synapses and some behavioral benefits in that model [38]. Beyond HD, correction of splicing deficits and consequent increases in the levels of survival motor neuron 2 protein in mouse models of spinal muscular atrophy improves hepatic phenotypes and greatly extends survival compared to treatments limited to the CNS [23]. Similarly, in a mouse model of the polyglutamine expansion disease spinal bulbar muscular atrophy, muscle expression of mutant androgen receptor was found to be required for disease-relevant phenotypes, excepting CNS protein aggregation [39]. Taken together with our results, it appears that the degree of cell autonomy for neurodegenerative disease phenotypes varies across both specific phenotypes and specific diseases, and is therefore worthy of careful study in each case.

We chose to test the links between peripheral organ pathology and CNS-resident signs of disease in the B6. *Htt^Q111/+^* knock-in mouse model of HD. Conducting interventional trials in this model, indeed any knock-in models of HD, is a minority position. In fact, a recent review suggests that only approximately 3% of preclinical studies in HD research are conducted using knock-in HD mice [40]. New investigations of knock-in models of HD reveal very clear, if subtle, behavioral changes [41,42], as well as extremely consistent molecular changes which are most pronounced in the striatum [16,43]. Power analyses suggest many of these phenotypes provide a robust system in which to test novel therapeutic compounds targeting the earliest phenotypes caused by CAG-expansion in *Htt*. Our own power analyses suggest that a multivariate suite of molecular phenotypes has excellent power to reveal successful reversal of these phenotypes in well-powered preclinical studies [15]. These mice enable us to examine therapies which may moderate the very earliest stages of HD, which precede synapse loss, gliosis or cell death. It is possible that some interventions may provide benefit for more distal disease states, which would be missed by our analysis. This focus on the earliest changes is a choice that necessarily excludes discovering interventions which are beneficial at later stages of disease.

In the present study we investigated behavioral, as well as pathological, signs of HD in the B6.*Htt^Q111/+^* mouse model. In addition to modest hypoactivity and thigmotaxis observed in an open field task, we investigated long-term explicit spatial memory using a novel object location task (Fig 4). These mice have previously been reported to show deficits in long-term (24 hour) but not short-term (15 minute) object recognition memory. In the present study, we do not observe any deficit in long-term novel object location memory in *Htt^Q111/+^* mice (data not shown), however we do observe an unexpected reduction in the total number of object investigations in 9-month-old *Htt^Q111/+^* mice compared to *Htt+/+* mice (Fig 4C). On average, *Htt^Q111/+^* mice investigated objects in their environment 19% fewer times than *Htt^+/+^* mice. Statistically, this is the most robust behavioral finding we have observed in *Htt^Q111/+^* at this age. A range of behavioral studies suggest that motivational changes reminiscent of apathy are early phenotypes of knock-in models of HD [44,42]. Prospective longitudinal study of HD mutation carriers reveal that apathy is unique among psychiatric symptoms in that it increases in severity across all stages of the disease, as well as being highly correlated with cognitive and motor impairment [45]. These findings suggest that our observations of reduced object investigations in *Htt^Q111/+^* mice could point to early motivational changes, a hypothesis we are continuing to investigate with more targeted behavioral tasks.

Our results reveal that peripheral huntingtin silencing, or treatment with this specific ASO, result in modest reductions in body weight by 10 months of age (Fig 5). We have here limited our investigations to the impact of peripheral huntingtin silencing on CNS-resident signs of disease, but new evidence suggests that complete silencing of huntingtin in adult animals leads to peripheral phenotypes, including unexpected fatal pancreatitis [46]. Conversely, overexpression of wild-type huntingtin leads to robust increases in both body weight and the size of a number of organs including heart, liver, kidneys and spleen, suggesting huntingtin levels in peripheral tissues may have important physiological impacts [47]. On-going investigations using tissues generated in the current study will focus on peripheral organ phenotypes caused by long-term huntingtin lowering.

## Methods

### Mice and genotyping

Female B6.*Htt^Q111/+^* (research resource identifier RRID:IMSR_JAX:003456) and wild-type littermates were acquired from the Jackson Laboratories (Bar Harbor, ME). This strain is congenic on the C57BL/6J background and its creation been described previously [14]. CAG tract lengths of *Htt^Q111/+^* mice ranged from 107-119 with an average of 114 (Table 1); no differences in CAG tract lengths existed between the treatment groups (F_(2, 49)_ = 0.98, p = 0.38). Upon arrival to Western Washington University (WWU) at 3 weeks of age, mice were housed in a partially reversed light cycle, lights on from 12 am to 12 pm, with *ad libitum* access to food and water. Genotype and treatment were balanced across cages. After habituating for 5 weeks, treatment began at 2 months (61 ± 2 days) of age and lasted for 8 months. Two interim silencing cohorts (n = 2 per arm, total N = 24) were established to verify HTT silencing prior to study completion at time points approximately one-third and two-thirds through the trial (at 11 and 22 weeks). All procedures were reviewed and approved by the animal care and use committee at WWU (protocol 14-006).

**Table 1.** CAG tract lengths of *Htt^Q111/+^* mice in the efficacy cohort.

| Treatment | Genotype | Measured CAG (SD) |
| --- | --- | --- |
| *Htt* ASO | *Htt*<sup>Q111/+</sup> | 114.5 (2.5) |
| Control ASO | *Htt*<sup>Q111/+</sup> | 114.1 (2.7) |
| Saline | *Htt*<sup>Q111/+</sup> | 113.3 (3.0) |

For genotyping, genomic DNA was extracted from 3-mm tail biopsies taken at weaning. Presence or absence of the mutant allele was determined by polymerase chain reaction (PCR) using the primers CAG1 (5-’ATGAAGGCCTTCGAGTCCCTCAAGTCCTTC-3’) [48] and HU3 (5’-GGCGGCTGAGGAAGCTGAGGA-3’) [49], as previously described. PCR products were separated by gel electrophoresis in 2.2% agarose DNA cassettes (Lonza) and visualized using the FlashGel System.

### Antisense Oligonucleotide Administration

Ionis Pharmaceuticals supplied *Htt*-targeting (Ionis 419637, ‘*Htt* ASO’) or off-target control (Ionis 141923, ‘control ASO’) ASOs, the latter with no sequence match in the mouse genome. Both ASOs were 20 nucleotide 5-10-5 2’-methoxyethyl gapmers with phosphorothioate backbones [17]. The sequences of *Htt* ASO and control ASO were 5’-<u>CCTGC</u>ATCAGCTTTA<u>TTTGT</u>-3’ and 5’-<u>CCTTC</u>CCTGAAGGTT<u>CCTCC</u>-3’ respectively (2’-methoxyethyl modified bases in the oligo ‘wings’ underlined). Mice were treated with 50 mpk *Htt* or control ASO via weekly IP injections, a dose and frequency selected based on initial dose-response and washout studies (Figure in S1-2 figures). As an additional treatment control, a smaller cohort of mice were treated with 4 μL/g body mass of saline.

### Tissue Harvesting

At 10 months of age, mice were sacrificed and tissues were harvested as described previously [15]. Briefly, a lethal injection of at least 250 mpk of sodium pentobarbital containing euthanasia solution was administered via IP injection, plasma was collected via cardiac puncture and centrifugation, and mice were transcardially perfused with phosphate buffered saline (PBS) to clear blood from organs. Terminal body mass, as well as liver, perigonadal white adipose tissue pads (WAT), interscapular brown adipose tissue (BAT), and spleen masses were recorded. Right lateral and caudate liver lobes were fixed overnight in 10% neutral buffered formalin (NBF) for histological analyses. To enable molecular analyses of peripheral tissues, the remaining liver tissue was flash frozen along with WAT, BAT, quadricep, and heart. Whole brain was removed and the hemispheres were separated along the longitudinal fissure. Using the Mouse Brain Slicer (Zivic), a consistent 3-mm corticostriatal block was cut from the left hemisphere and fixed overnight in NBF. Striatum, cortex, and cerebellum were dissected from the right hemisphere and flash frozen for molecular analyses.

### MSD Assay

BioFocus, a Charles River company (Leiden, The Netherlands) quantified levels of both mutant and total HTT using previously described MSD assays [29]. Each assay utilized rabbit polyclonal pAB146 as the capture antibody, but differing sets of secondary antibodies in order to discriminate between polyglutamine expanded HTT and total HTT. Tissue lysates were divided and incubated in either the mouse monoclonal MW1 antibody (Developmental Studies Hybridoma Bank; Ab_528290) for mutant HTT assays or the rabbit polyclonal pAB137 antibody and the monoclonal mouse anti-HTT antibody (EMD Millipore: MAB2166; Ab_2123255) for total HTT assays. A goat anti-mouse SULFO TAG antibody was used to detect all secondary antibodies. HTT quantification was performed using flash frozen liver, WAT and BAT tissue from a subset of five *Htt^+/+^* and *Htt^Q111/+^* mice per arm of the efficacy trial. Likewise, hemi-striatal, liver, WAT and BAT from two mice per treatment and genotype condition in the interim silencing cohorts were used to assess total and mHTT knockdown mid-way through the efficacy trial.

### Behavior

Open field and novel object location testing occurred in an open field arena constructed of black acrylic walls with a white acrylic floor (44 × 44 × 44 cm). Behavioral experiments were carried out between 1:00 and 8:00 PM each day, and mice were moved into the experimental room 30 minutes prior to each session to habituate. Room lighting was maintained at approximately 475 lux during all behavioral assays. Between trials the floor, walls of the arena, and objects were cleaned thoroughly using 70% EtOH to minimize the effect of olfactory cues on exploratory behavior. On days 1 and 2, mice were habituated to the open-field arena in the absence of any objects and permitted to explore freely for 10 minutes before being returned to their home cage. Activity was tracked using an overhead camera, and the exploratory behavior recorded during day 1 was analyzed using Noldus Ethovision XT 8 [50].

The novel object location task was conducted on day 3, as described [51]. Briefly, mice were placed into the open-field arena in the presence of two identical objects (Erlenmeyer flasks, terra cotta pots, Nalgene bottles filled with blue food coloring, and Nalgene bottles covered with textured coozies) located in the NW and NE corners of the arena. Objects were counterbalanced between animals to ensure equal representation between experimental conditions. Large intra-maze cues (a square, a triangle, an equal sign, and a plus) were taped to the N, S, W, and E walls respectively to provide spatial cues. Mice were placed into the apparatus facing N toward the objects and were permitted to explore for a single 10 minute acquisition period before being returned to their home cage. Following a 24 hour delay, one of the objects was displaced to the diametrically opposite corner of the arena (i.e. if the NW object was displaced, it was moved to the SW corner, if the NE object was displaced, it was moved to the SE corner). Mice were placed into the open-field facing N and permitted to explore for a single 5 minute probe session. The number of explorations toward each object, defined as orienting the head toward the object within a 2 cm proximity or interacting directly with the object, was recorded by an experimenter blind to genotype and treatment.

### Transcriptional Profiling

Total RNA was extracted from the right hemistriatum as described previously [15] for both RNA sequencing (RNAseq) and quantitative reverse-transcription polymerase chain reaction (QRT-PCR). RNAseq was conducted at EA | Q^*2*^ Solutions with cDNA libraries prepared using the Illumina TruSeq Stranded mRNA sample preparation kit (Illumina # RS-122-2103). Quality control as well as gene and isoform quantification were performed with an EA | Q^*2*^ Solutions developed analysis pipeline, mRNA v7. Bowtie version 0.12.9 was used to align reads to the mouse transcriptome, followed by quantification with RSEM version 1.1.18.

In preparation for QRT-PCR, messenger RNA was reverse transcribed using the Superscript III First Strand Synthesis System (Life Technologies) according to the manufacturer’s protocol. QRT-PCR was conducted and analyzed as described [15] using the following taqman probes purchased from Life Technologies: Drd1a: Mm02620146\_s1, N4bp2: Mm01208882\_m1, Islr2: Mm00623260\_s1, and Pde10a: Mm00449329\_m1. All transcripts were normalized to β-actin.

### Immunohistochemistry

Fixed corticostriatal blocks were paraffin embedded, cut into 5-μm sections, and mounted on glass slides for immunohistochemistry at Histology Consultation Services (Everson, WA). Experimenters blind to treatment conducted deparaffinization, p62 staining, imaging and analysis as described previously [15].

### Statistical Analyses

All data were analyzed using R version 3.3.1 [52]. Data presented in figures 1,2,3b,4, S3 and S5 were tested for normality (Shapiro-Wilk test) and homoscedasticity (Levene’s test). All data met or approximated parametric assumptions within groups and were fit with linear models analysed by ANOVA. Data were presented as boxplots created with ggplot2 [53] - horizontal lines indicate 25th, 50th and 75th percentile, while the vertical whiskers indicate the range of data. Data falling outside 1.5 times the interquartile range are graphed as isolated points, but were not excluded from statistical analysis. RNASeq data were analyzed using the limma [54] package of Bioconductor [55]. Body weight data was analyzed using a minP-based parametric bootstrap multiple comparison procedure [36,37].

## Supplemental Figures

**S1 Fig.**
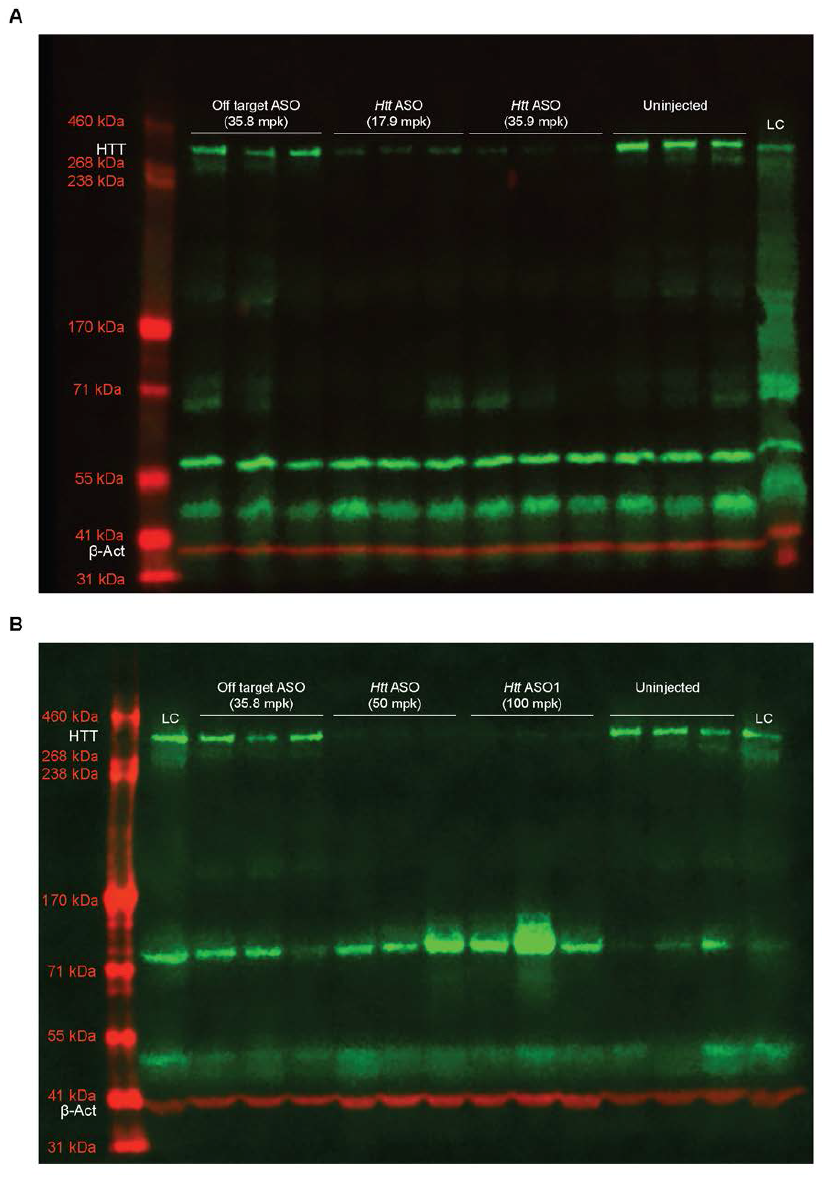
Treatment with 50 mpk or greater of *Htt* ASO effectively suppresses HTT in the liver. In order to determine an appropriate dose of *Htt* ASO, we conducted a preliminary dose response study and quantified HTT levels in the liver (our primary tissue of interest) via western blotting. Due to gel constraints, all doses could not be included on a single gel, therefore 17.9 and 35.8 mpk are presented in (A) while 50 and 100 mpk are presented in (B). Faint HTT bands remain in liver protein extracted from mice treated with 35.8 mpk per week (A), while treatment with 50 mpk per week or greater produced seemingly complete knockdown of HTT (B). Accordingly, 50 mpk was selected as the *Htt* ASO dose for the efficacy study. Abbreviations - LC: positive loading control, HTT: huntingtin protein, β-Act: β-Actin

**S2 Fig.**
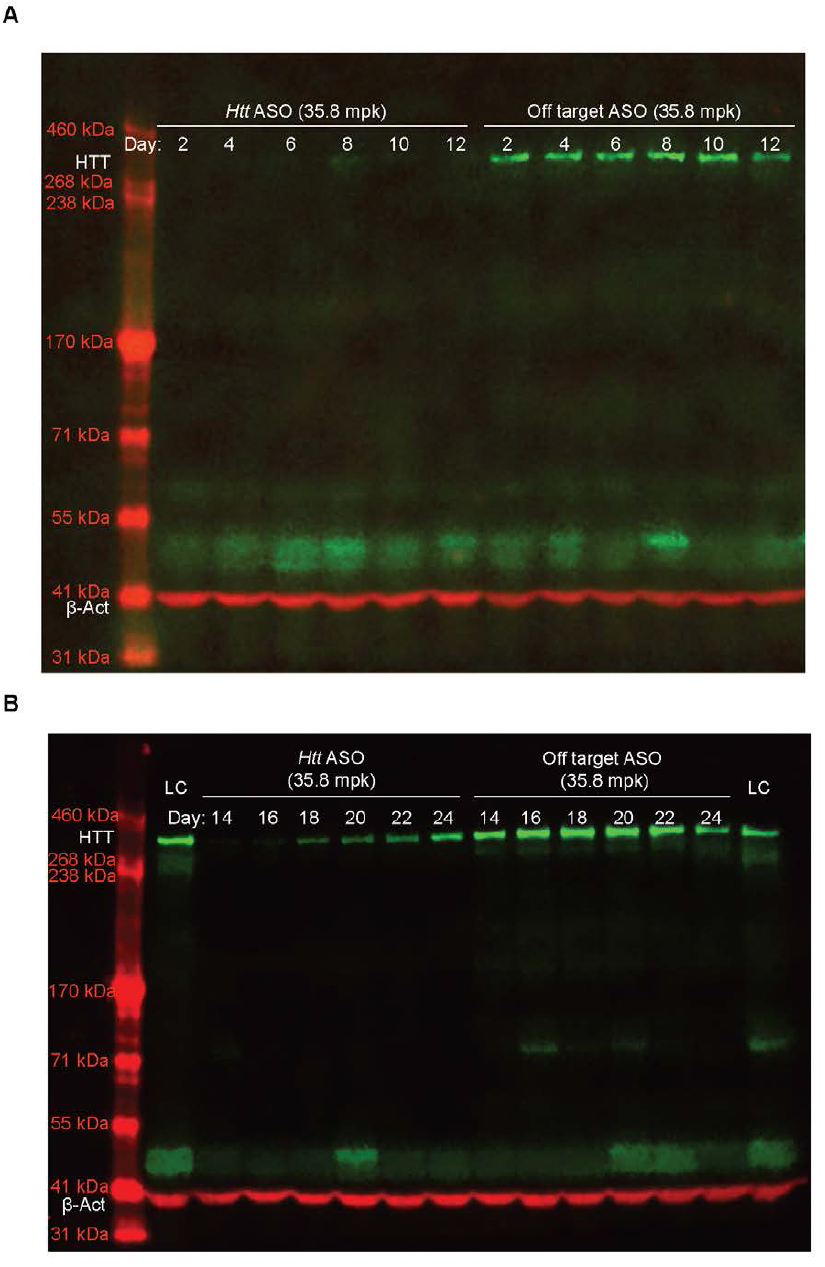
*Htt* ASO mediated suppression lasts for 14 days after cessation of treatment in the liver. To roughly characterize the duration of action of our chosen ASO, we performed three, weekly IP injections of *Htt* ASO or off target ASO and measured liver HTT levels every other day for 24 days. Due to gel constraints, samples could not be loaded on a single gel, therefore days 2 - 12 are shown in (A) and days 14 - 24 are shown in (B). Based on these observations, we concluded weekly IP injections of *Htt* ASO were sufficient to ensure no recovery of HTT levels between treatments. Abbreviations - LC: positive loading control, HTT: huntingtin protein, β-Act: β-Actin

**S3 Fig.**
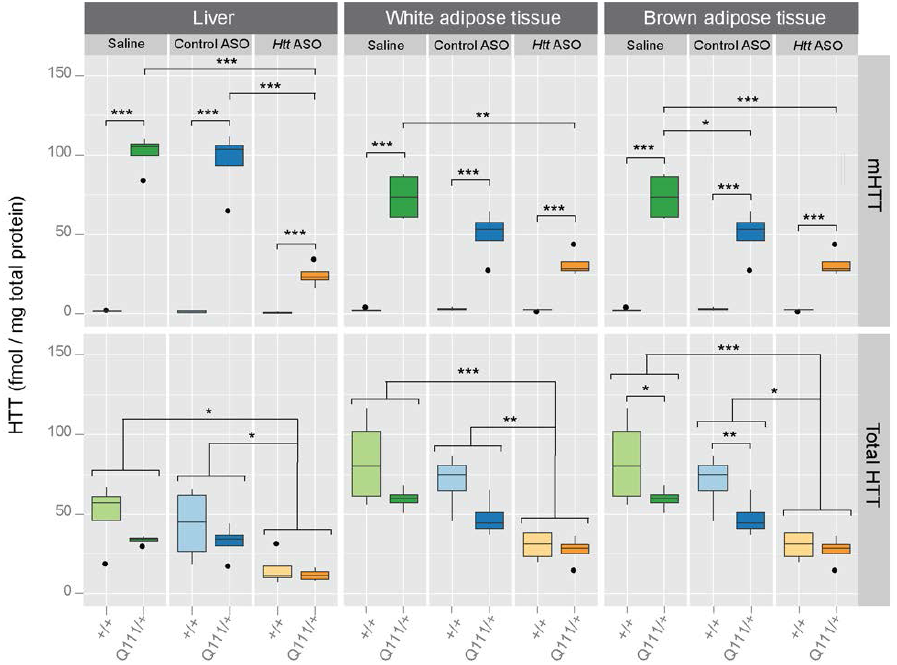
ASO-mediated HTT suppression confirmed at intermediate timepoints throughout the efficacy trial, suggesting continuous HTT knockdown. Both total and mHTT levels were quantified via MSD assay in tissues harvested from two interim silencing cohorts at 4.5- and 7-months of age. As expected, the extent and pattern of HTT suppression was consistent with that seen in the efficacy trial, suggesting lapses in HTT knockdown during the efficacy trial are unlikely. Not surprisingly, mHTT levels were observed to be significantly higher in *Htt^Q111/+^* mice than *Htt^+/+^* mice across tissues. HTT levels following *Htt* ASO treatment were significantly reduced compared to saline or control treatment.* p ≤ 0.05, ** p ≤ 0.01, *** p ≤ 0.001: by Tukey’s HSD pairwise comparisons

**S4 Fig.**
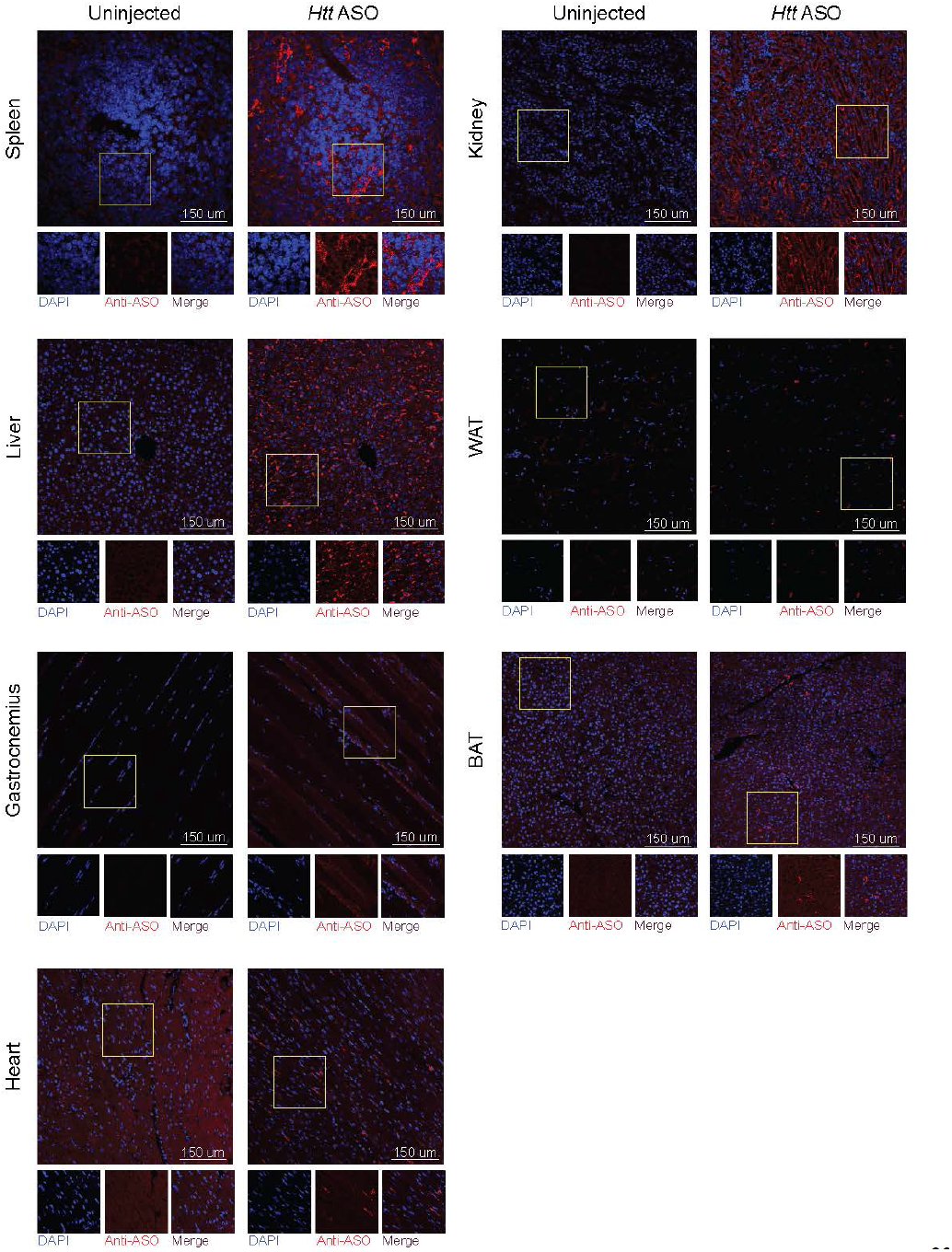
Qualitative assessment of peripheral *Htt* ASO distribution reveals efficient targeting of liver, spleen, and kidney. After 1 month of treatment with 50 mpk *Htt* ASO per week, mice were sacrificed and *Htt* ASO uptake was evaluated using an antibody reactive to the ASO backbone. Out of seven peripheral tissues, Htt ASO uptake was most pronounced in the liver, kidney and spleen, with modest uptake evident in the perigonadal white adipose tissue, gastrocnemius, interscapular brown adipose tissue and heart.

**S5 Fig.**
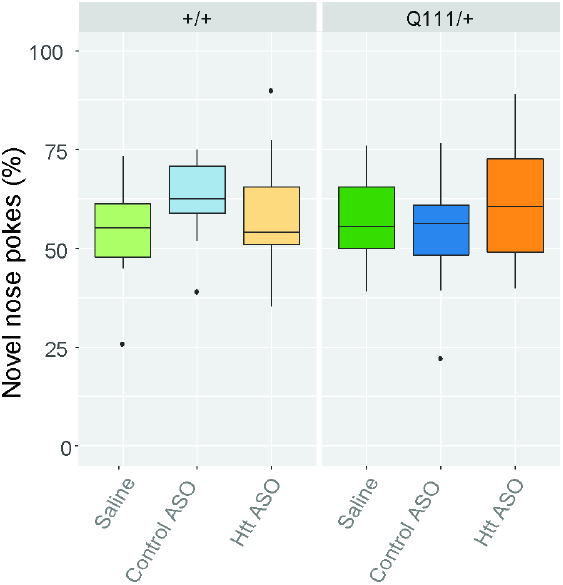
Regardless of treatment, *Htt^Q111/+^* and *Htt^+/+^* mice prefer objects in novel locations over familiar locations. In every genotype and treatment condition, mice explored objects located in the novel location over 50% of the time, demonstrating that spatial long term memory is neither impaired in *Htt^Q111/+^* mice or affected by *Htt* ASO treatment.

## Supplemental Results and Methods

### Results

#### Dose-response and washout studies

We piloted four different doses of *Htt* ASO and qualitatively assessed HTT knockdown by western blotting. At the two highest doses, no HTT could be detected, despite robust HTT signal in protein lysates extracted from off target ASO and uninjected mice (Figure in S1 Figure). Therefore we selected 50 mg per kg body mass (mpk) per week as the dose for the efficacy study; this was the minimal dose at which complete HTT knockdown was observed. β-Actin was used as a loading control and was successfully detected in all samples regardless of treatment.

To determine how frequently *Htt* ASO must be administered in order to maintain constant HTT knockdown, we conducted a washout study. Wild type mice were given weekly injections of *Htt* ASO or off target ASO for three weeks and sacrificed every other day for 24 days. During this time, we observed complete HTT silencing for 14 days after cessation of treatment (Figure in S2 Figure). Even 24 days after the final injection, HTT levels did not recover to physiological levels. For the efficacy trial, we chose to conservatively administer *Htt* ASO via weekly injections.

#### HTT knockdown in interim silencing cohorts

At two points midway through the efficacy trial, we confirmed HTT knockdown in three peripheral tissues of interest: liver, perigonadal white adipose tissue, and interscapular brown adipose tissue. HTT levels were quantified via mesoscale discovery assays, as described in the primary manuscript. No differences in mutant or total HTT suppression patterns were observed between timepoints, therefore data were grouped for statistical analysis. Htt ASO treatment had a significant effect on total HTT levels (Figure in S3 Figure) in all three tissues (effect of treatment in the liver: *F*_(2,18)_ = 9.8, p = 1.8 × 10^−3^, white adipose tissue: *F*_(2, 18)_ = 15.7, p = 1.2 × 10^−4^, and brown adipose tissue: *F*_(2, 18)_= 11.3, *p* = 6.5 × 10^−4^). As expected, Tukey’s HSD *post hoc* comparisons revealed *Htt* ASO treatment suppressed HTT compared to both control ASO (liver: *p* = 8.8 × 10^−3^, white adipose tissue: *p* = 3.1 × 10^×3^, and brown adipose tissue: *p* = 2.2 × 10^−3^) and saline treatment (liver: *p* = 2.6 × 10^−3^, white adipose tissue: *p* = 1 × 10^−4^, and brown adipose tissue: *p* = 1.4 × 10^−3^). Similarly, mHTT was significantly reduced by *Htt* ASO treatment in all tissues quantified (effect of treatment in the liver: *F* _(2,18)_ = 36.8, *p* = 4.4 × 10^−7^, white adipose tissue: *F* _(2,18)_ = 9.6, *p* = 1.4 × 10^−3^, and brown adipose tissue: *F*_(2,18)_= 5.6, *p* = 0.01). Tukey’s HSD *post hoc* tests verified reduction of mHTT following *Htt* ASO treatment compared to saline treatment (liver: *p* = 1.2 × 10^−6^, white adipose tissue: *p* = 1 × 10^−3^, and brown adipose tissue: *p* = 0.01). Interestingly, no difference in mHTT levels was observed by Tukey’s HSD *post hoc* comparisons between *Htt* ASO and control ASO treatment in either of the adipose tissues (white adipose tissue: *p* = 0.16, and brown adipose tissue: *p* = 0.11), despite *Htt* ASO treatment reducing mHTT levels relative to control treatment in the liver (*p* = 3.5 × 10^−6^).

### Methods

#### Mice

In order to determine an appropriate ASO dose and frequency of administration, we conducted dose-response (n = 5 per arm, total N = 30) and washout studies (n = 1 per arm, total N = 24) in male, wild-type C57BL/6J mice (stock no. 000664) acquired from the Jackson Laboratories (Bar Harbor, ME). Mice were housed under standard vivarium conditions with *ad libitum* access to food and water. After shipment, mice habituated to local vivarium conditions for at least two weeks prior to every study. For both the washout and dose-response studies, treatment began at three months of age and continued for three weeks. For the dose-response study, mice were sacrificed two days after the final injection as described in the primary manuscript. For the washout study, mice were sacrificed every other day after cessation of treatment.

ASO uptake in peripheral tissues was qualitatively assessed in small, tissue distribution cohort made up of female, C57BL/6J mice bred locally (n = 2 per arm, total N = 4). Mice in the tissue distribution cohort were maintained with *ad libitum* access to food and water. Treatment began at 4 months (121 ± 1 days) of age and continued for four weeks. At 5 months (147 ± 4 days) of age, or two days after cessation of treatment, mice were sacrificed via intraperitoneal injection of at least 250 mpk sodium pentobarbital and transcardially perfused with phosphate buffered saline (PBS). Liver, perigonadal white adipose tissue (WAT), interscapular brown adipose tissue (BAT), kidney, spleen, heart, and gastrocnemius were dissected and fixed overnight in 10% neutral buffered formalin. All procedures were reviewed and approved by the animal care and use committee at Western Washington University (protocol 14-006).

### Antisense Oligonucleotide Administration

In an initial dose-response study, we administered a pan-*Htt* targeted ASO (Ionis 419637, *Htt* ASO) via weekly IP injection at four different doses: 17.9, 35.8, 50, or 100 mpk. As a control, an off target ASO was administered weekly at 35.8 mpk. The off target ASO was selected as a control because it shares a similar chemical structure with *Htt* ASO, both of which are 20 nucleotide 5-10-5 2’-methoxyethyl gapmers. In addition, we compared both *Htt* ASO and off target ASO treated mice to uninjected mice. Based on this initial dose-response study, we selected a dose of 50 mpk per week for all subsequent studies.

To roughly characterize *Htt* ASO duration of action, we conducted a washout study in which mice were treated with *Htt* ASO or off target ASO at 35.8 mpk per week for three weeks. After cessation of treatment, mice were sacrificed every other day for 24 days and HTT levels quantified by western blot.

For the tissue distribution study, mice received weekly injections of *Htt* ASO at 50 mpk for four weeks.

### Immunoblotting

Tissue (half of dissected brown adipose tissue and 35 - 55 mg of liver tissue) was transferred to BeadBug tubes prefilled with 1.5-mm zirconium oxide beads (Benchmark) and homogenized for 2 minutes at 4,000 rpm using the BeadBug Microtube Homogenizer (Benchmark). Protein was extracted using 450 μL of RIPA buffer (150 mM NaCl, 25 mM Tris-HCl, 1% NP-40, 1% sodium deoxycholate, 0.1% SDS) containing protease and phosphatase inhibitors (Thermo Fisher Scientific). Concentration of protein lysates were determined by Pierce BCA assay (Thermo Fisher Scientific) according to the manufacturer’s protocol.

Total HTT levels were quantified via western blotting. Equal amounts of protein were loaded into 3-8% Tris-Acetate Mini Gels (Thermo Fisher Scientific) and separated electrophoretically by molecular weight. Protein was transferred at 4–℃20 for hours17 to a PVDF membrane.

Western blot assay membranes were blocked for 1 hour at room temperature using Odyssey Blocking Buffer (LiCor) before incubating for 1 hour at room temperature or overnight at 4 antibodies. Monoclonal primary antibodies were diluted 1:1,000 in Odyssey blocking buffer plus 0.2% tween and consisted of mouse anti-HTT (EMD Millipore: MAB2166; Ab_2123255) and rabbit anti-β-Actin (Cell Signalling: mAB#8457, Ab_10950489). Membranes were rinsed three times in PBS plus 0.05% tween (PBST), then incubated in secondary antibodies for 1 hour at room temperature. Polyclonal secondary antibodies were IRDye 800CW conjugated goat anti-mouse IgG (LiCor: 926- 32210; Ab_621842) and IRDye 680RD conjugated goat anti-rabbit IgG (LiCor: 926-68071; Ab_10956166) diluted 1:15,000 in Odyssey blocking buffer plus 0.2% tween and 0.01% SDS. After rinsing three times in PBST, membranes were washed in PBS before imaging for 2 minutes each at 700 and 800 nm.

### Immunohistochemistry and Imaging

As part of the tissue distribution study, formalin-fixed liver, WAT, BAT, gastrocnemius, heart, spleen, and kidney tissues were paraffin embedded, cut into 5-μm sections, and mounted on glass slides. Sections were deparaffinized and heat mediated antigen retrieval was conducted in Tris-EDTA buffer before staining overnight in primary anti-ASO antibody (1:1,000) at 4 in goat anti-rabbit secondary antibody conjugated to Alexa568 (1:1,000) for one hour at room temperature and mounted with DAPI fluoromount-G (Southern Biotech). All confocal images were acquired with a IX-81 laser-scanning confocal microscope using Fluoview 1000 software (Olympus). Image processing and analysis was performed as described [1].

